# Discrimination against non-nestmates functions to exclude socially parasitic conspecifics in an ant

**DOI:** 10.1101/2024.05.16.593084

**Authors:** Takuma P. Nakamura, Shigeto Dobata

## Abstract

Social animals utilise various communication methods to organise their societies. In social insects, nestmate discrimination plays a crucial role in regulating colony membership. Counter to this system, socially parasitic species employ diverse behavioural and chemical strategies to bypass their host’s detection. In this study, we tested whether such parasitic adaptations could be detected in the incipient stage of social parasitism that is observed as intraspecific phenomena in some social insects. The Japanese parthenogenetic ant *Pristomyrmex punctatus* harbours a genetically distinct cheater lineage which infiltrates and exploits host colonies. We found that intrusion of this intraspecific social parasite was defended by nestmate discrimination of host colonies without any behavioural strategies specialised in social parasitism. Most of the cheaters were eliminated through aggressions by host workers that are typically observed against non-nestmates, resulting in a low intrusion success rate for the cheaters (6.7%). This result contrasts with the expectation from interspecific social parasitism but rather resembles the intraspecific counterpart reported in Cape honeybees (*Apis mellifera capensis*), illustrating the role of nestmate discrimination against the intrusion of intraspecific social parasites.

## 1 INTRODUCTION

Social insects, such as ants, bees, wasps and termites, employ a diverse range of communication methods to organise their societies (Wilson, 1971; 1975). Nestmate recognition, a pivotal faculty among these social insects, entails the identification of individuals either as integral members of their colony or as extraneous entities, thereby regulating the admission of individuals into the colony (van Zweden & d’Ettorre, 2010). The recognition typically relies on chemical cues that are specific to each colony (Breed *et al*., 1995; Gamboa *et al*., 1986; Morel *et al*. 1988). Especially in ants, body cuticular hydrocarbons (CHCs) are primary cues in nestmate recognition (Bonavita-Cougourdan *et al*., 1987; Lahav *et al*., 1999; Sturgis & Gordon, 2012). Individuals producing distinct CHC blends are identified as non-nestmates (Guerrieri *et al*., 2009; Ozaki *et al*., 2005). After nestmate recognition, agonistic defensive behaviours are directed toward non-nestmate intruders, known as nestmate discrimination (Richard & Hunt, 2013).

Nestmate recognition/discrimination is regarded as an adaptive colony defence against the intrusion of heterospecific and non-nestmate conspecific individuals (Crozier & Pamilo, 1996; Hölldobler & Wilson, 1990). These intruders may reap fitness benefits by exploiting the socially acquired resources of the colonies. Some species, called social parasites, have evolved to specialise in such socially parasitic strategies (Cini *et al*., 2019). These strategies include xenobiosis, in which the parasite cares for its own brood but cannot live without the host; temporary parasitism, in which the parasite depends on the host’s workforce only during the founding phase of the parasite’s colony; dulosis, in which the parasite enslaves workers of host colonies; and inquilinism, in which the parasite usually tolerates the host queen but entirely depends on the host’s workforce for reproduction (Buschinger, 2009).

The social parasites typically use chemical and behavioural strategies to avoid detection by the host individuals and thereby facilitating their invasion (Cervo, 2006; Huang & Dornhaus, 2008; Lenoir *et al*., 2001). For example, some social parasites reduce chemical odour of their body surface to make them less likely to be detected and attacked by their hosts (Dronnet *et al*., 2005; Lorenzi & Bagnères, 2003; Nehring, 2014). In socially parasitic ants with chemical camouflage, the parasites actively perform allogrooming toward host individuals which allows them to gather chemical cues from the host body surface (Lenoir *et al*., 2001). All these strategies can be primarily assessed by behavioural observations as an initial step in the study of sophisticated social parasitism.

The elaboration of social parasitism depends on the evolutionary history of host-social parasite relationships. Social parasites, especially inquilines, are often the closest relatives of their host species, known as Emery’s rule (Huang & Dornhaus, 2008 and references therein). The parasitic strategies devised by social parasites in their incipient stages would shed light on how the social parasitism evolves and contributes to the speciation from the host (Bourke & Franks, 1991; Savolainen & Vepsäläinen, 2003). In this study, we explored the existence of an infiltration strategy in a socially parasitic “cheater” lineage of the Japanese parthenogenetic ant, *Pristomyrmex punctatus*.

The life history of *P. punctatus* is highly unique, distinguished by the absence of the queen caste, the scarcity of males, and the annual life cycle (Itow *et al*., 1984; Tsuji & Dobata, 2011). Most females are workers in their external morphology while producing offspring with two ovarioles through thelytokous parthenogenesis. Their reproductive division of labour is rather based on adult age, known as age-polyethism, where young adults undertake inside-nest tasks, laying eggs and nurturing the brood, while older adults cease reproduction and participate in outside-nest tasks such as foraging (Tsuji, 1988b, 1990b). The outside-nest workers often engage in mutualistic associations with lycaenid butterflies by exchanging defence and nutrition, which is known to be facilitated by the recognition of specific CHCs (Hojo *et al*., 2014; Hojo *et al*., 2015).

Previous studies have found the “cheater” lineage from a local population in central Japan (Sasaki & Tsuji, 2003; Dobata *et al*., 2009; Dobata *et al*., 2011; Dobata & Tsuji 2013; Tsuji & Dobata 2011). The cheaters exhibit genetically determined morphology distinct from the typical workers (i.e., their hosts), such as distinctively larger body size and four ovarioles, while they are parthenogenetic and annual as well (Tsuji & Dobata 2011). They coexist with typical workers in a colony but are infrequently active outside the nest, refrain from nursing or foraging, and exhibit higher egg-laying rates compared to typical workers (Sasaki & Tsuji, 2003; Dobata & Tsuji 2013). The increased proportion of the cheaters in a colony causes labour shortage and results in the collapse of the entire colony (Dobata & Tsuji 2013). Phylogenetic analyses demonstrated that the cheater lineage is monophyletic and is nested within worker lineages of *P. punctatus*, indicating that this social parasitism is an intraspecific phenomenon (Dobata *et al*., 2009; 2011).

A previous population genetic study (Dobata *et al*., 2011) showed that the cheaters with identical genotypes were found in genetically diverse host colonies, the degree of which was not likely to occur only by colony fission and thus suggested that the cheaters migrate among colonies with low frequencies, despite the operational nestmate discrimination system (Sanada-Morimura *et al*., 2003; Tsuji, 1988a). This finding renders the cheaters in *P. punctatus* analogous to interspecific social parasites, or more precisely inquilines in some ants.

We anticipated that the cheater lineage devises any specialised behavioural strategies to circumvent nestmate discrimination, which could be assessed by observing their behaviours during intrusion and settlement into host colonies. For example, lowered host aggression toward the cheaters compared with the other non-nestmates would suggest the existence of chemical mimicry or stealth. Likewise, peaceful survival of the cheaters inside host nest sites would indicate their successful intrusion. Alternatively, given the detrimental impact of cheaters on the fitness of host workers, the host workers might have elaborated counter-adaptation that enables them to detect and selectively eliminate the cheaters. In either case the host-intruder behavioural interactions involving the cheaters should differ from those involving non-nestmate workers. Through the implementation of a colony introduction experimental design and video recording, we systematically explored the existence of such strategies and evaluated the success of cheater intrusion after 24 hours of introduction.

## 2 MATERIALS AND METHODS

### 2.1 Sampling

A total of 38 colonies of *P. punctatus* were excavated in Kihoku, Mie Prefecture, Japan, from May 18 to 20, 2022. Individuals of the cheater lineage have distinctly larger body size compared with those of worker lineages, and the resulting bimodal distribution of body size among nestmates (Dobata *et al*., 2009) practically enabled visual identification of cheaters. The presence/absence of putative cheaters was confirmed in the field by spreading the colony on a white plastic tray and visually inspecting individual body sizes for 30 person-minutes. Consequently, 14 colonies (6 with and 8 without putative cheaters) were brought back to the laboratory, were housed in plastic trays containing nest material (soil and dead leaves) and were subjected to natural light-dark cycles at 25°C. Glass test tubes were employed as nesting sites, filled with water and covered with absorbent cotton. To prevent ant escape, the tray walls were coated with Fluon®. The colonies received insect jelly (Pro Jelly, KB Farm, Japan) twice a week. The presence of cheaters was finally confirmed by picking up the larger individuals and checking the presence of three conspicuous ocelli on the head (Tsuji & Dobata, 2011). The colonies eventually used in the experimental procedures are listed in Table S1.

### 2.2 Colony introductions

We conducted a “colony introduction” assay following the methodology outlined by Roulston *et al*. (2003). We used five field-collected colonies lacking the cheaters for the source of hosts (A, B, C, D, and F; hereafter referred to as host-source colonies) and five containing the cheaters for the source of intruders (H, I, J, K, and M; intruder-source colonies). The experimental flow is shown in Figure 1: in each trial, we established an experimental host colony composed of 150 workers which were haphazardly taken from a host-source colony, taking advantage of queenlessness of *P. punctatus*. Subsequently, an individual, either cheater, nurse, or forager, was picked up from an intruder-source colony, which was then introduced to the host colony as a non-nestmate intruder to observe her interactions with host workers and her survival. The 15 pairs of host-source and intruder-source colonies tested are listed in Table S2. As a negative control, a nestmate nurse from the host-source colony was introduced into the experimental host colony, which was repeated three times to match the number of trials with those for a host-intruder pair. Nurses are generally expected to be less aggressive than foragers (Larsen *et al*. 2016), which is true for *P. punctatus* (T. Nakamura and S. Dobata, personal observation), and we deemed them appropriate for the negative control. The experiments were conducted from August 22 to September 11, 2022, during a later stage of brood production.

**Figure 1.**
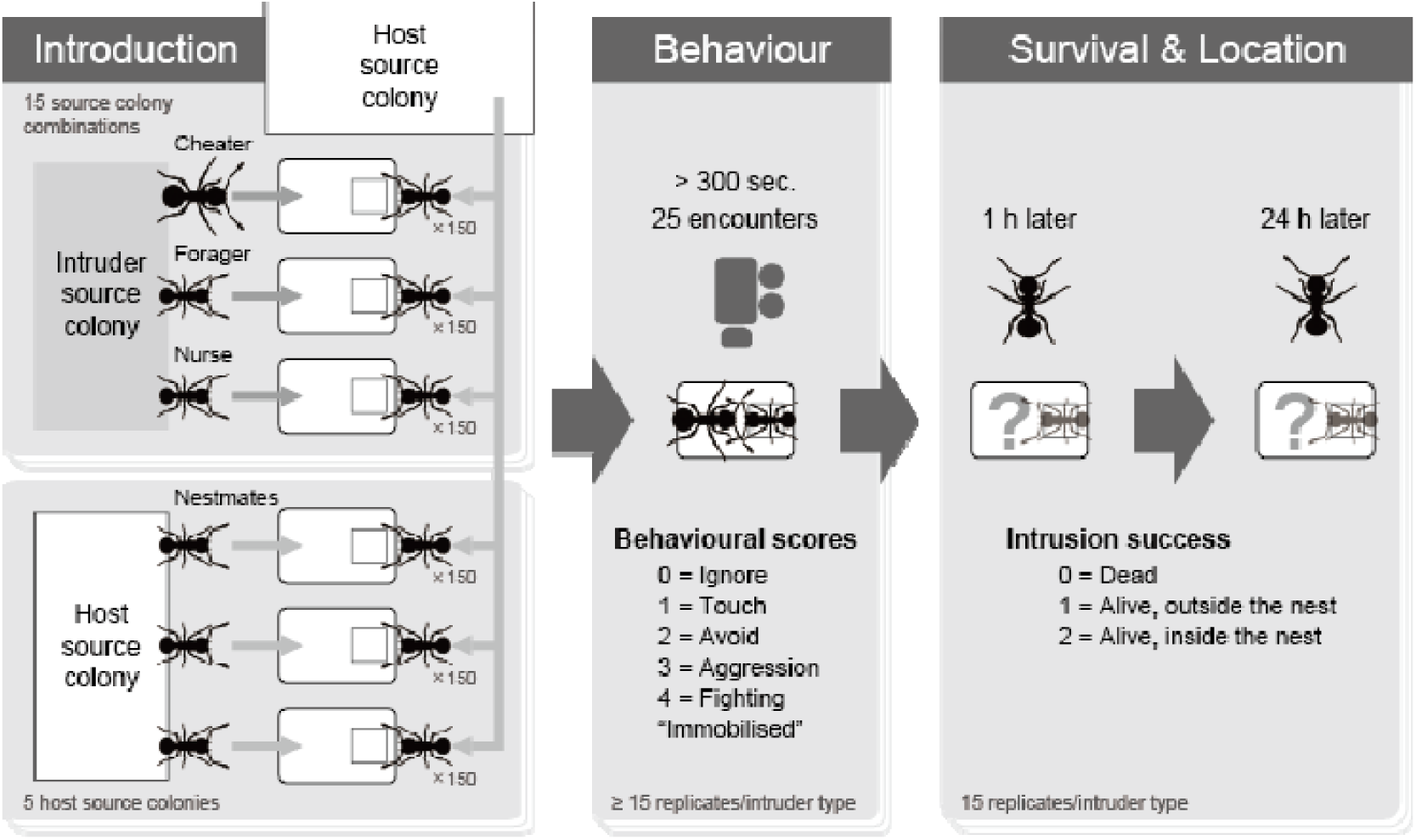
Outline of the experimental flow.

The detailed procedure of a trial is described below: Plastic containers (120 mmL × 80 mmW × 70 mmH) with a plaster floor and Fluon®-coated walls were prepared. Additionally, a shallow square hole (approximately 27 mm square, 2 mm depth) was created on the floor as a nest site, covered with a 30 mm square glass plate (Figure 2). A total of 150 adults, including nurses and foragers, were haphazardly taken from a host-source colony and were placed in a container. Most individuals became distributed in the nest site within 24 hours (Figure 2) and thereby formed a host colony. The holes in the cases were moistened by allowing water to seep through the surrounding plaster.

**Figure 2.**
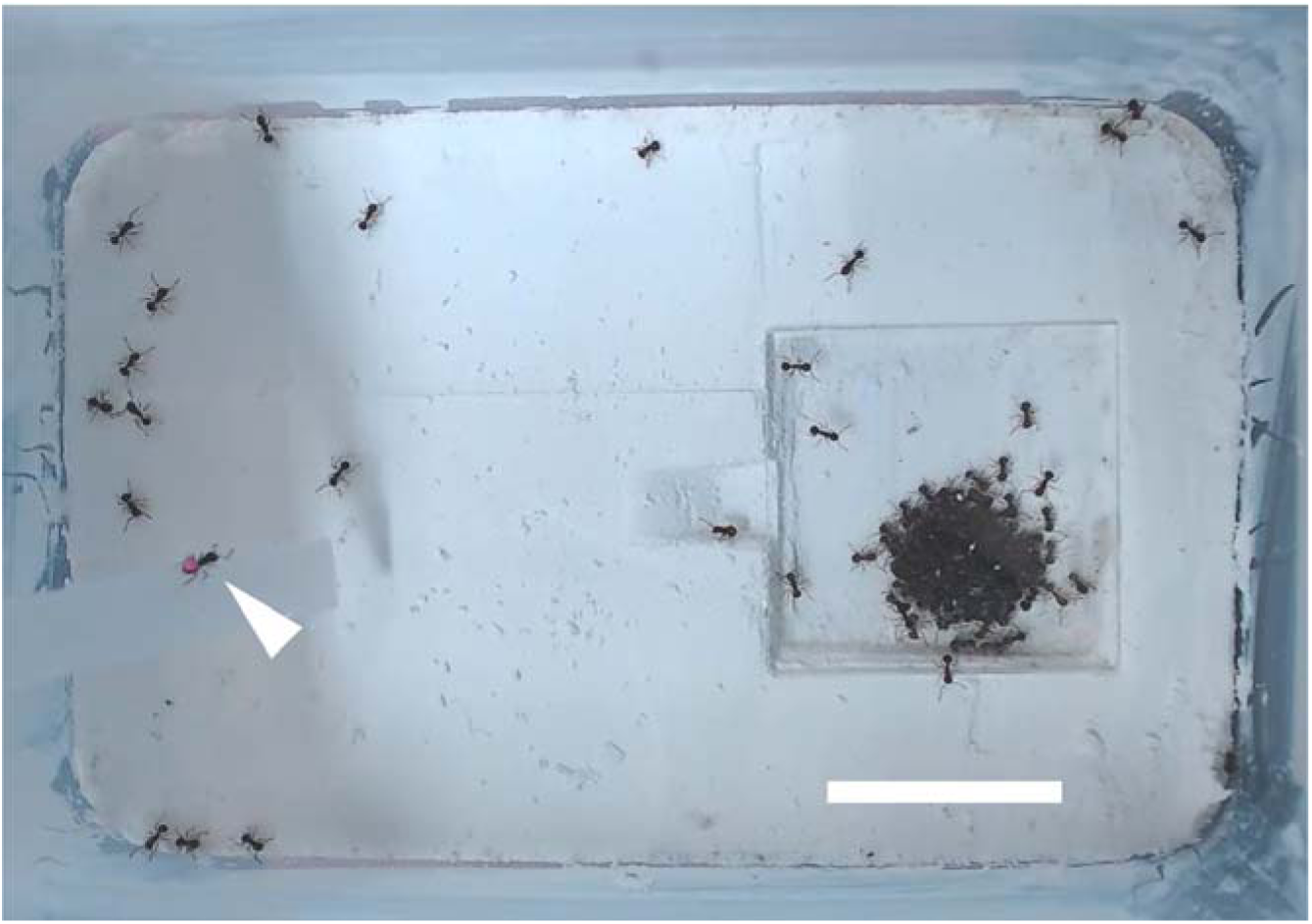
An experimental host colony with an intruder (arrowhead) being introduced. Scale bar: 20 mm.

On the same day, intruders were extracted from their source colony as follows: the test tube was first removed from the tray and placed in a plastic container; cheaters were individually picked up initially based on their body size and were finally confirmed by checking the presence of the three ocelli as described above; the test tube were then maintained in the container for an hour and individuals outside the test tube were separated every hour as foragers, repeated for three times which enables the removal of ca. 80% of foragers from the mixture of workers (Tsuji 1994); and finally to confirm the behavioural status directly, individuals were forced to exit the test tube and those who carry broods (eggs, larvae, or pupae) were separated as nurses. In case no broods were available in the nest, broods from another intruder-source colony were temporarily given to confirm their behaviours as follows: in each test trial a brood was picked up from the test tube and was presented in front of the test workers. The worker who held the brood in her mandibles was regarded as a nurse. The brood was then swiftly rescued within ca. 30-60 seconds. The separated individuals were marked on the abdomen with a paint marker (Paint Marker px-20 Pink, Mitsubishi Pencil, Tokyo, Japan) and were placed in another container with a sufficient number (ca. 1000) of individuals from their source colony until used in the introduction assays on the next day.

Approximately 24–26 hours after the host colony was established, a single painted individual was introduced onto the foraging arena of the host colony using a plastic bridge (Figure 2). The entire container was video-recorded with a Logicool C920n HD PRO webcam and Logicool Capture software (Logicool Co., Tokyo, Japan), which started just before the introduction and lasted for at least 300 consecutive seconds. All recordings were performed between 1000 and 1800 hours. One hour after introduction, we checked inside the container to observe the survival and location (inside or outside of the nest) of the intruder. Individuals unable to move were tentatively treated as dead, after confirming that they still did not move when picked up by the experimenter with tweezers. The nest site was then moistened again as described above. After 24 hours of introduction, the same checking procedure was conducted. The intruder was then collected and stored at –80°C. Sometimes the paint mark was removed from the intruder after introduction, preventing identification of nurses and foragers and resulting in missing data in some trials (*n* = 1 for foragers after 1 h; *n* = 4 for nurses and *n* = 3 for foragers after 24 h). In such cases, we repeated the above procedure using the same combination of host-source and intruder-source colonies with the same intruder type up to three times (detailed in Table S2). Nevertheless, to avoid the potential survivorship bias caused by preferential selection of intruders retaining the paint mark, these additional replicates were eventually included only in the behavioural analysis where all the intruders retained the paint mark (described in 2.3), while they were omitted from the analyses of survival and location of the intruders.

### 2.3 Behavioural analysis

We analysed the video-recorded behaviours of the intruder and the host workers during the colony introductions. Each video was assigned a random ID to blind the intruder types as much as possible. Behavioural scores were derived from Suarez *et al*. (1999) with slight modification: 0 = ignore (a focal individual shows no interest even though the interactant makes contact using her antennae); 1 = touch (antennal contacts); 2 = avoid (an encounter resulting in the retreat of a focal individual); 3 = aggression (a focal individual opens their mandibles to threaten or bite the interactant); 4 = fighting (prolonged biting, often resulting in the immobilisation of the interactant); and an additional category “immobilised”. These scores were assigned for the behaviours of the intruder and a host worker, respectively, at each encounter. An encounter was recorded when at least one individual showed actions scored as 1 or higher. We recorded the behavioural scores until 25 encounters were made by an intruder, following the method of Roulston *et al*. (2003). In the case of an encounter consisting of more than one behavioural type (e.g., 1 and 3), it was represented by the behaviour with the highest score.

### 2.4 Statistical analyses

All statistical tests were conducted using R 4.3.2 (R Core Team, 2023) with the significance level α = 0.05. The effects of intruder types on the degree of hostility at each encounter were examined through the application of mixed-effect logistic regressions. The datasets of intruders and host workers were analysed separately. The model formulae in Wilkinson notation were

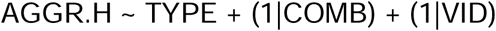

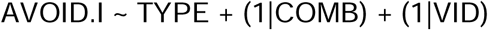

Based on functionality of behaviours, we analysed the host’s aggressive behaviour (scores 3 and 4 combined) and the intruder’s avoiding behaviour (score 2). The proportion of these behaviours out of (25 – counts of “immobilised”) encounters were treated as response variables (AGGR.H and AVOID.I, respectively). The types of intruders (TYPE) were treated as the fixed effect. Identifiers for the combination of source colonies (COMB) and those of the videos (VID) were treated as random effects (random intercepts). In addition, the proportion of the outcome “immobilised” out of 25 encounters was similarly analysed. Model fitting was executed using the glmer() function from the “lme4” package. The effect of intruder type was examined using the likelihood ratio test (LRT) implemented in the anova() function, by dropping TYPE from the full model. Post-hoc pairwise comparisons among the four intruder types were conducted using the Tukey method, implemented through the glht() function in the “multcomp” package.

We next focused on behaviours of non-nestmate intruders (cheaters, foragers and nurses) and analysed how they co-occurred with those of host workers at each encounter, using the mixed-effect logistic regressions:

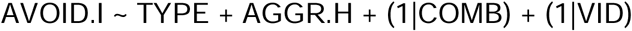

The response variable AVOID.I was the occurrence of the intruder’s avoiding behaviour (score 2) at each encounter, and the explanatory variables were the intruder type (TYPE) and the corresponding occurrence of the aggressive behaviour (scores 3 and 4 combined) of the host worker (AGGR.H). The observations in which either was “immobilised” were removed from this analysis, and the other procedures were the same as described above. An additional analysis was performed to assess the effects of the above explanatory variables on the intruder’s response “immobilised” at each encounter. The procedure was the same as above except for the inclusion of observations in which either was “immobilised”.

For the fate of the intruders, the data were treated as a variable FATE with an ordinal scale: level 0 (dead), level 1 (alive outside the nest) and level 2 (alive inside the nest). Only the first introduction trial was considered for each non-nestmate type × source colony combination (see 2.2), resulting in 15 replicates per intruder type. To examine whether the intruder fate was explained by the intruder type (TYPE), the mixed-effect logistic regression (1 hour; there were only two levels 1 and 2) or mixed-effect ordinal logistic regression (24 hours) models were fitted to the data, with the latter model implemented in the clmm() function of the package “ordinal”:

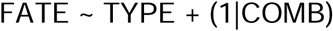

Identifiers for the combination of source colonies (COMB) were treated as a random effect (random intercept). The effect of TYPE was examined using LRT with anova() or anova.clm() functions. The post-hoc pairwise comparisons among intruder types were conducted using the Tukey method, implemented in glht() function of the “multcomp” package for the logistic regression and functions in the “emmeans” packages for the ordinal logistic regression.

## 3 RESULTS

### 3.1 Behavioural analysis

The composition and correspondence of behavioural scores observed among intruders and host workers are visually summarised in Figure 3 and Figure S1. Among host workers, aggressive behaviours (i.e., scores 3 and 4 out of the observations except “immobilised”) were more frequently observed when invaded by non-nestmates than by nestmates, and the difference was statistically significant (logistic regression; LRT, χ^2^ = 12.425, d.f. = 3, *p* = 0.0061; multiple comparisons with Tukey method; |*z*| ≥ 2.988, *p* ≤ 0.0132; Figure 4A).

**Figure 3.**
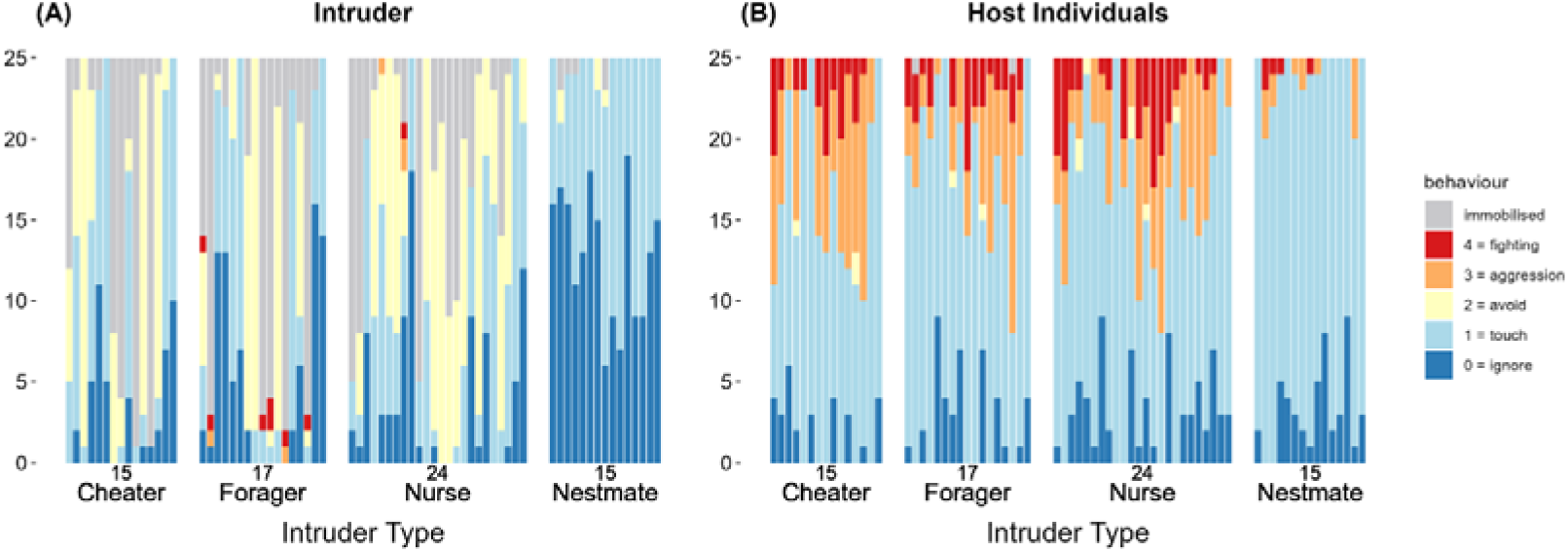
Frequencies of behavioural categories exhibited by (A) intruders and (B) host workers in colony introduction experiments. Each vertical bar corresponds to an introduction trial, with their left-to-right arrangement indicating the same trial between (A) and (B).

**Figure 4.**
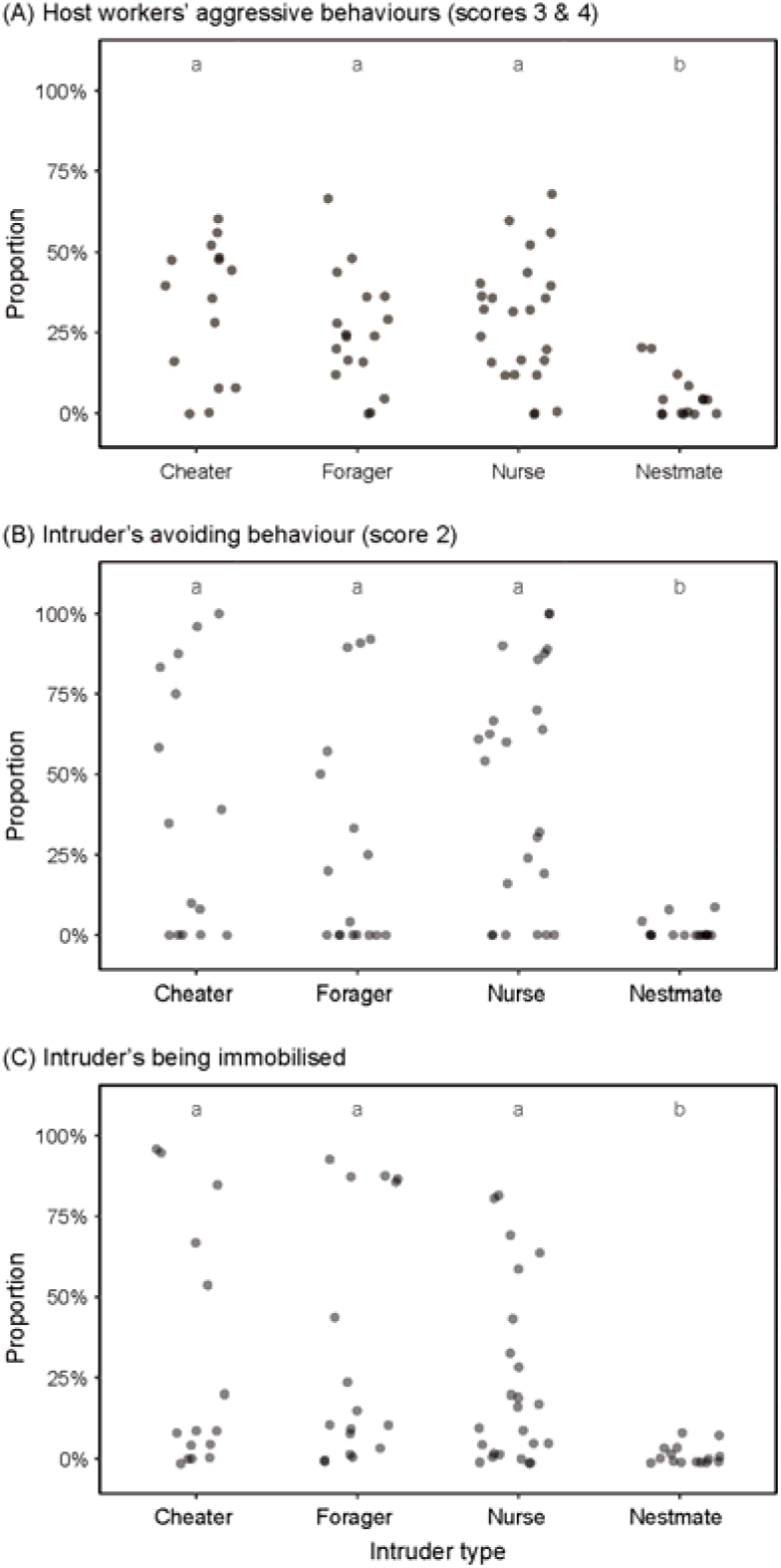
Proportions of specific behaviours exhibited by interactants, comparing among the four intruder types. (A) Aggressive behaviours (scores 3 and 4) by host workers, (B) Avoiding behaviours (score 2) by intruders, and (C) immobilisation of intruders by the host workers. Different letters at the top indicate statistically significant differences among intruder types. In (A) and (B), the encounters that resulted in “immobilised” were omitted from the denominators.

Meanwhile, the avoiding behaviour (i.e., the proportion of behavioural score = 2) was more frequently exhibited by non-nestmate intruders than by nestmate intruders, and the difference was again statistically significant (logistic regression; LRT, χ^2^ = 19.826, d.f. = 3, *p* = 0.0002; multiple comparisons with Tukey method; |*z*| ≥ 2.941, *p* ≤ 0.0164; Figure 4B). Likewise, non-nestmate intruders were more likely to be immobilised than nestmate intruders (logistic regression; LRT, χ^2^= 10.647, d.f. = 3, *p* = 0.0138; multiple comparisons with Tukey method; |*z*| ≥ 2.582, *p* ≤ 0.0452; Figure 4C).

In the interactions between host workers and non-nestmate intruders, statistically significant positive associations were detected between the host’s aggressive and the intruder’s avoiding behaviours (logistic regression; slope ± SE = 1.5868 ± 0.2614; LRT, χ^2^ = 41.365, d.f. = 1, *p* < 0.0001), and between the host’s aggressive behaviour and the intruder’s immobilisation (logistic regression; slope ± SE = 1.4455 ± 0.1910; LRT, χ^2^ = 57.715, d.f. = 1, *p* < 0.0001).

Meanwhile, the effect of non-nestmate intruder type was neither statistically significant for the intruder’s avoiding behaviour (LRT, χ^2^ = 4.5334, d.f. = 2, *p* = 0.1037), nor for the intruder’s immobilisation (LRT, χ^2^ = 2.2676, d.f. = 2, *p* = 0.3218). An additional analysis showed that adding the interaction term between the non-nestmate intruder type and the host’s aggressive behaviour (i.e., TYPE:AGGR.H) was not statistically significant both for the intruder’s avoidance (LRT, χ^2^ = 2.1198, d.f. = 2, *p* = 0.3465) and for the intruder’s immobilisation (LRT, χ^2^ = 5.7937, d.f. = 2, *p* = 0.0552).

### 3.2 Survival and location of intruders

After an hour of introduction, all intruders survived while non-nestmate intruders were less likely to be found inside the nest than nestmate nurses, to a statistically significant degree (logistic regression; LRT, χ^2^ = 18.429, d.f. = 3, *p* = 0.0004; multiple comparisons with Tukey method; |*z*| ≥ 2.544, *p* ≤ 0.0364) (Figure 5A). These results indicate lower rates of early success for non-nestmates in intruding into the host’s nest. After 24 hours, all nurses, irrespective of their source colony of origin (non-nestmate or nestmate), exhibited continued survival, while non-nestmate foragers and cheaters showed lowered survival rates (Figure 5B). Using the ordinary scale of the intruder’s success as a response, the nestmate and non-nestmate nurses showed greater success in intruding into the host’s nest than non-nestmate foragers and cheaters (ordinal logistic regression; LRT, χ^2^ = 41.707, d.f. = 3, *p* < 0.0001; multiple comparisons with Tukey method; |*z*| ≥ 3.076, *p* ≤ 0.0113). One out of 15 non-nestmate cheaters (6.7%) survived for 24 hours and successfully intruded into the host’s nest site.

**Figure 5.**
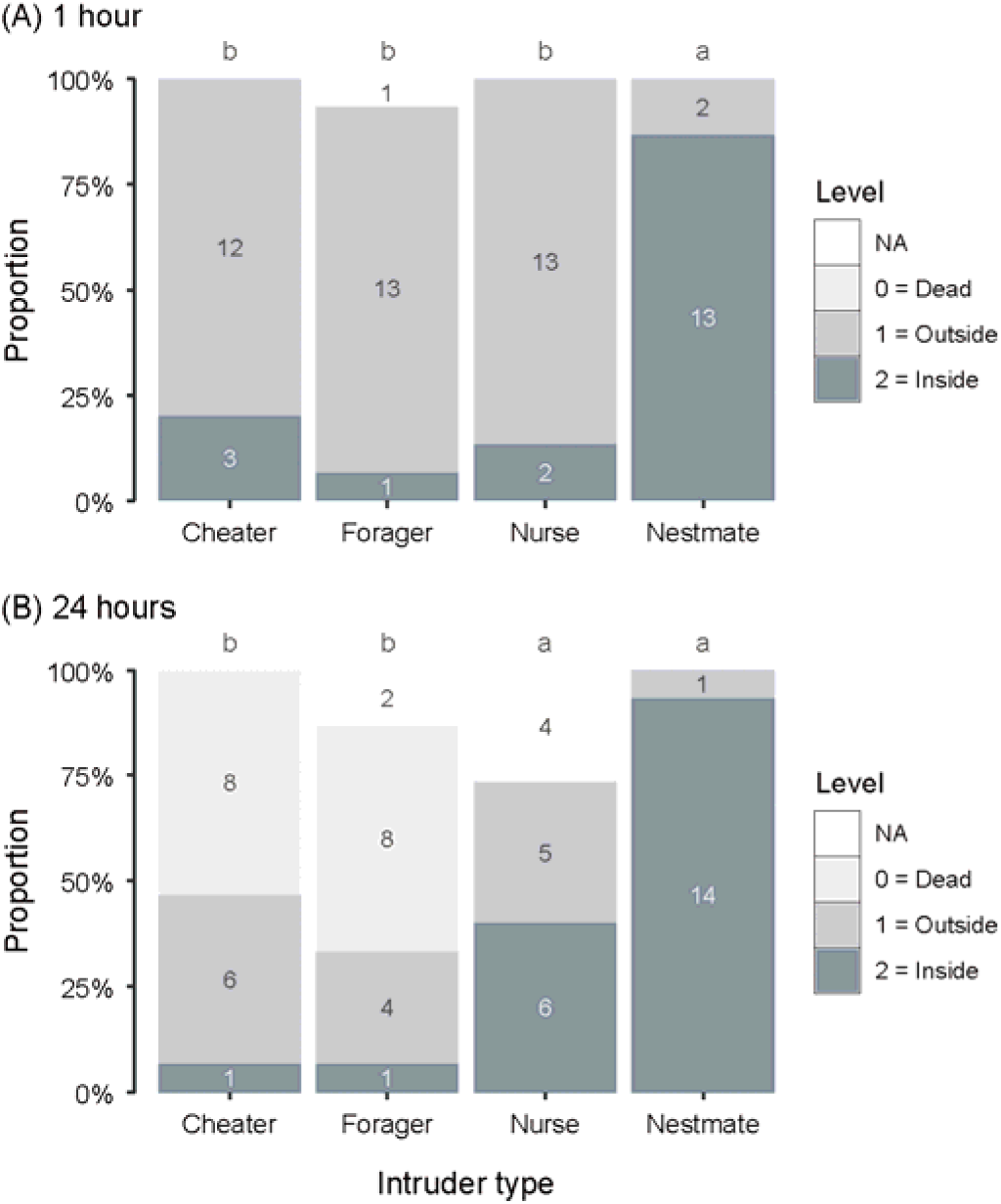
Results of colony introduction experiments. (A) after 1 hour and (B) after 24 hours of introduction. Different letters at the top indicate statistically significant differences among intruder types. Numbers in the bars indicate the number of individuals in each state.

## 4 DISCUSSION

In general, evolved social parasites are expected to show adaptive traits that help them to reap benefits from their hosts. We aimed at identifying such adaptation at the initial stage of host-social parasite interactions, namely the intrusion into the host colony. A previous population genetic study suggested migration of cheaters among host colonies (Dobata et al., 2011). Therefore, our initial hypothesis presumed that cheaters would develop behavioural strategies to evade nestmate discrimination.

Nonetheless, we did not observe any distinct behaviours in the cheaters which contribute to social parasitism. No statistically significant differences were detected between cheaters and the other non-nestmate intruders in the frequencies of avoiding behaviours, in the frequencies of being immobilised by host workers, nor in the manner of reactions to host workers’ aggressive behaviours (i.e., the interaction term between the non-nestmate intruder type and the host’s aggressive behaviour). These results provide support against the presence of adaptive behaviours of cheaters when infiltrating other colonies. Our analyses also suggested that aggressive behaviours of host workers against the intruders did not differ between cheaters and the other non-nestmate intruders. These findings suggest that the cheaters are treated simply as non-nestmates by host workers, at least in the intrusion stage. In this respect, cheaters of *P. punctatus* contrast with other socially parasitic ants that employ chemically distinctive strategies to evade detection and aggression by hosts, such as mimicry, camouflage and stealth (Akino, 2008; Lenoir *et al*., 2001; Nehring *et al*., 2015).

The apparent lack of any intrusion strategies might be explained by the relatively short evolutionary history of the cheater lineage in *P. punctatus*, which was previously estimated as 200–9200 generations (Dobata *et al*., 2011). In other words, sufficient generation time has not yet elapsed for the cheaters to adapt against the nestmate discrimination. Nonetheless, direct evidence for the lack of chemical strategies in the cheaters awaits further study examining CHCs in *P. punctatus*. Likewise, our study does not rule out the possibility of any post-intrusion strategies, behavioural or chemical, which become functional after the successful settlement of cheaters in a new host colony (Cini *et al*., 2019). A previous study found that a higher proportion of cheaters in a colony was associated with a higher tendency for a nestmate worker to be found outside their nest (i.e., to be a forager; Dobata & Tsuji, 2013). Future studies should examine whether any manipulative and counter-manipulative features can characterise the cheater-worker interactions in *P. punctatus*.

The cheaters showed lowered success rates of intrusion compared with nestmates (Figure 5), which is a typical outcome of ant workers in the face of non-nestmate colonies. In the present experiment, the probability with which a cheater successfully settled in the host’s nest after 24 hours was 6.7% (= 1/15; Figure 5B). Given that the intrusion of the cheaters leads to the eventual collapse of the host colonies within several generations, how the observed intrusion success contributes to the persistence of the social parasite lineages deserves further theoretical considerations (cf. Brandt *et al*., 2005). It is worth noting that a higher probability of intrusion success would act as a “double-edged sword” by leading to the pandemic-like exhaustion of the entire host population.

A similar but slightly higher probability of intrusion success (15%) was observed in the Cape honeybee, *Apis mellifera capensis* (Beekman *et al*., 2002). In this honeybee subspecies, female workers can reproduce through thelytokous parthenogenesis, invade colonies of the subspecies *A. m. scutellata*, engage in social parasitism, and ultimately dismantle the host colony (Neumann & Moritz, 2002; Yagound *et al*., 2020). However, they lack a distinct strategy during intrusion and are often rejected by the host *A. m. scutellata* workers (Neumann & Pirk, 2019).

Both in *P. punctatus* and in *A. m. capensis*, it can be argued that the nestmate discrimination serves as a border defence against intrusion of the conspecific and exogenous cheaters. In general, nestmate discrimination functions as defence mechanisms against any selfishness exhibited by non-nestmates, including food-theft, brood-robbing and brood parasitism (Breed *et al*., 2012; Downs & Ratnieks, 2000), and against pathogens likely attached to non-nestmates (Lemanski *et al*., 2021). These two species offer promising systems to study how the nestmate discrimination has been evolutionarily shaped against the invasion of intraspecific social parasites, which could be understood in parallel with the evolution of immune systems in multicellular organisms against cancer cells (Dobata & Tsuji, 2009; Dujon *et al*., 2020; Pull & McMahon, 2020).

An unexpected result of our experiment was that all non-nestmate nurses survived after 24 hours of experimental introduction, with over a half found inside the host nest-site (Figure 5B). This result was in stark contrast with the fate of non-nestmate foragers and cheaters, which showed lowered survival as expected from non-nestmate discrimination. Despite encountering aggression and rejection upon introduction, the non-nestmate nurses appeared to gain acceptance from their host colony. In *P. punctatus*, nurses are younger adults and are capable of thelytokous reproduction, whereas foragers cease reproduction with degenerated ovarioles (Tsuji, 1990b). Therefore, from the kin selection point of view, the apparent selective acceptance of non-nestmate nurses should lower the relatedness within a host colony and thus diminish the inclusive fitness of the host worker. Previous studies found a certain level of flexibility in the nestmate discrimination in *P. punctatus* (Sanada-Morimura *et al*., 2003; Tsuji, 1990a). Although further examination is beyond the scope of our current study, future work is needed to clarify the fate of non-nestmate nurses as a potential source of genetic heterogeneity observed in the field colonies (Nishide *et al*., 2007; Satow *et al*., 2013).

## ACKNOWLEDGEMENTS

We thank Kazuki Tsuji, Hiroyuki Shimoji, Tetsu Morimoto, Po-Wei Hsu, Norika Kaneda, and Haruka Asanabe for their help during the field and laboratory work. This work was partly supported by Grant-in-Aid for Scientific Research (KAKENHI; 21H02537, 21H04885, 22H02705, and 22H02364 to SD) from the Japan Society for the Promotion of Science.

## Notes

https://doi.org/10.5061/dryad.59zw3r2h2

### Competing Interest Statement

The authors have declared no competing interest.

### Summary of Updates

The entire manuscript was substantially rewritten, including the title change. Information about the study system was added in the introduction; Study design was clarified in Figure 1; statistical analyses and their visualization were totally revised to address mixed effect models.

## REFERENCES

1. Akino, T. (2008). Chemical strategies to deal with ants: A review of mimicry, camouflage, propaganda, and phytomimesis by ants (Hymenoptera: Formicidae) and other arthropods. Myrmecological News, 11, 173–181.

2. Beekman, M., Wossler, T. C., Martin, S. J., & Ratnieks, F. L. W. (2002). Parasitic Cape honey bee workers (*Apis mellifera capensis*) are not given differential treatment by African guards (*A. m. scutellata*). Insectes Sociaux, 49(3), 216–220. doi:10.1007/s00040-002-8304-0

3. Bonavita-Cougourdan, A., Clement, J. L. & Lange, C. (1987). Nestmate recognition: the role of cuticular hydrocarbons in the ant *Camponotus vagus* Scop. Journal of Entomological Science, 22, 1–10. doi:10.18474/0749-8004-22.1.1

4. Bourke, A. F. G., & Franks, N. R. (1991). Alternative adaptations, sympatric speciation and the evolution of parasitic, inquiline ants. Biological Journal of the Linnean Society, 43(3), 157–178. doi:10.1111/j.1095-8312.1991.tb00591.x

5. Brandt, M., Foitzik, S., Fischer-Blass, B., & Heinze, J. (2005). The coevolutionary dynamics of obligate ant social parasite systems–between prudence and antagonism. Biological Reviews, 80(2), 251–267. doi:10.1017/S1464793104006669

6. Breed, M. D., Garry, M. F., Pearce, A. N., Hibbard, B. E., Bjostad, L. B., & Page, R. E. (1995). The role of wax comb in honey bee nestmate recognition. Animal Behaviour, 50(2), 489–496. doi:10.1006/anbe.1995.0263

7. Breed, M. D., Cook, C., & Krasnec, M. O. (2012). Cleptobiosis in social Insects. Psyche: A Journal of Entomology, 484765. doi:10.1155/2012/484765

8. Buschinger, A. (2009). Social parasitism among ants: a review (Hymenoptera: Formicidae). Myrmecological News, 12(3), 219–235.

9. Cervo, R. (2006). *Polistes* wasps and their social parasites: An overview. Annales Zoologici Fennici, 43, 531–549.

10. Cini, A., Sumner, S., & Cervo, R. (2019). Inquiline social parasites as tools to unlock the secrets of insect sociality. Philosophical Transactions of the Royal Society B: Biological Sciences, 374(1769), 20180193. doi:10.1098/rstb.2018.0193

11. Crozier, R. H., & Pamilo, P. (1996). Evolution of Social Insect Colonies: Sex Allocation and Kin Selection. Oxford University Press.

12. Dobata, S., Sasaki, T., Mori, H., Hasegawa, E., Shimada, M., & Tsuji, K. (2009). Cheater genotypes in the parthenogenetic ant *Pristomyrmex punctatus*. Proceedings of the Royal Society B: Biological Sciences, 276(1656), 567–574. doi:10.1098/rspb.2008.1215

13. Dobata, S., Sasaki, T., Mori, H., Hasegawa, E., Shimada, M., & Tsuji, K. (2011). Persistence of the single lineage of transmissible “social cancer” in an asexual ant. Molecular Ecology, 20(3), 441–455. doi:10.1111/j.1365-294X.2010.04954.x

14. Dobata, S., & Tsuji, K. (2009). A cheater lineage in a social insect. Communicative & Integrative Biology, 2(2), 67–70. doi:10.4161/cib.7466

15. Dobata, S., & Tsuji, K. (2013). Public goods dilemma in asexual ant societies. Proceedings of the National Academy of Sciences, 110(40), 16056–16060. doi:10.1073/pnas.1309010110

16. Downs, S. G., & Ratnieks, F. L. W. (2000). Adaptive shifts in honey bee (Apis mellifera L.) guarding behavior support predictions of the acceptance threshold model. Behavioral Ecology, 11(3), 326–333. doi:10.1093/beheco/11.3.326

17. Dronnet, S., Simon, X., Verhaeghe, J.-C., Rasmont, P., & Errard, C. (2005). Bumblebee inquilinism in *Bombus* (*Fernaldaepsithyrus*) *sylvestris* (Hymenoptera, Apidae): Behavioural and chemical analyses of host-parasite interactions. Apidologie, 36(1), 59–70. doi:10.1051/apido:2004070

18. Dujon, A. M., Gatenby, R. A., Bramwell, G., MacDonald, N., Dohrmann, E., Raven, N., Schultz, A., Hamede, R., Gérard, A.-L., Giraudeau, M., Thomas, F., & Ujvari, B. (2020). Transmissible cancers in an evolutionary perspective. iScience, 23(7), 101269. doi:10.1016/j.isci.2020.101269

19. Gamboa, G. J., Reeve, H. K., Ferguson, I. D., & Wacker, T. L. (1986). Nestmate recognition in social wasps: The origin and acquisition of recognition odours. Animal Behaviour, 34(3), 685–695. doi:10.1016/S0003-3472(86)80053-7

20. Guerrieri, F. J., Nehring, V., Jørgensen, C. G., Nielsen, J., Galizia, C. G., & d’Ettorre, P. (2009). Ants recognize foes and not friends. Proceedings of the Royal Society B: Biological Sciences, 276(1666), 2461–2468. doi:10.1098/rspb.2008.1860

21. Hojo, M. K., Pierce, N. E., & Tsuji, K. (2015). Lycaenid caterpillar secretions manipulate attendant ant behavior. Current Biology, 25(17), 2260–2264. doi:10.1016/j.cub.2015.07.016

22. Hojo, M. K., Yamamoto, A., Akino, T., Tsuji, K., & Yamaoka, R. (2014). Ants use partner specific odors to learn to recognize a mutualistic partner. PLoS one, 9(1), e86054. doi:10.1371/journal.pone.0086054

23. Hölldobler, B., & Wilson, E. O. (1990). The Ants. Harvard University Press.

24. Huang, M. H., & Dornhaus, A. (2008). A meta-analysis of ant social parasitism: Host characteristics of different parasitism types and a test of Emery’s rule. Ecological Entomology, 33(5), 589–596. doi: 10.1111/j.1365-2311.2008.01005.x

25. Itow, T., Kobayashi, K., Kubota, M., Ogata, K., Imai, H. T., & Crozier, R. H. (1984). The reproductive cycle of the queenless ant *Pristomyrmex pungens*. Insectes Sociaux, 31(1), 87–102. doi:10.1007/BF02223694

26. Lahav, S., Soroker, V., Hefetz, A., & Vander Meer, R. K. (1999). Direct behavioral evidence for hydrocarbons as ant recognition discriminators. Naturwissenschaften, 86(5), 246–249. doi:10.1007/s001140050609

27. Larsen, J., Nehring, V., d’Ettorre, P., & Bos, N. (2016). Task specialization influences nestmate recognition ability in ants. Behavioral Ecology and Sociobiology, 70, 1433–1440. doi:10.1007/s00265-016-2152-9

28. Lemanski, N., Silk, M., Fefferman, N., & Udiani, O. (2021). How territoriality reduces disease transmission among social insect colonies. Behavioral Ecology and Sociobiology, 75(12), 164. doi:10.1007/s00265-021-03095-0

29. Lenoir, A., D’Ettorre, P., Errard, C., & Hefetz, A. (2001). Chemical ecology and social parasitism in ants. Annual Review of Entomology, 46, 573–599. doi:10.1146/annurev.ento.46.1.573

30. Lorenzi, M. C., & Bagnères, A. G. (2002). Concealing identity and mimicking hosts: A dual chemical strategy for a single social parasite? (*Polistes atrimandibularis, Hymenoptera: Vespidae*). Parasitology, 125(6), 507–512. doi:10.1017/S003118200200238X

31. Morel, L., Vander Meer, R. K., & Lavine, B. K. (1988). Ontogeny of nestmate recognition cues in the red carpenter ant (*Camponotus floridanus*). Behavioral Ecology and Sociobiology, 22(3), 175–183. doi:10.1007/BF00300567

32. Nehring, V., Dani, F. R., Turillazzi, S., Boomsma, J. J., & d’Ettorre, P. (2015). Integration strategies of a leaf-cutting ant social parasite. Animal Behaviour, 108, 55–65. doi:10.1016/j.anbehav.2015.07.009

33. Neumann, P., & Moritz, R. (2002). The Cape honeybee phenomenon: The sympatric evolution of a social parasite in real time? Behavioral Ecology and Sociobiology, 52(4), 271–281. doi:10.1007/s00265-002-0518-7

34. Neumann, P., & Pirk, C. W. W. (2019). Increased response to sequential infections of honeybee, Apis mellifera scutellata, colonies by socially parasitic Cape honeybee, A. m. capensis, workers. Scientific Reports, 9(1), 7582. doi:10.1038/s41598-019-43920-1

35. Nishide, Y., Satoh, T., Hiraoka, T., Obara, Y., & Iwabuchi, K. (2007). Clonal structure affects the assembling behavior in the Japanese queenless ant *Pristomyrmex punctatus*. Naturwissenschaften, 94(10), 865–869. doi:10.1007/s00114-007-0267-6

36. Ozaki, M., Wada-Katsumata, A., Fujikawa, K., Iwasaki, M., Yokohari, F., Satoji, Y., Nisimura, T., & Yamaoka, R. (2005). Ant nestmate and non-nestmate discrimination by a chemosensory sensillum. Science, 309(5732), 311–314. doi:10.1126/science.1105244

37. Pull, C. D., & McMahon, D. P. (2020). Superorganism Immunity: A Major Transition in Immune System Evolution. Frontiers in Ecology and Evolution, 8. doi:10.3389/fevo.2020.00186

38. R Core Team (2023). R: A language and environment for statistical computing. R Foundation for Statistical Computing, Vienna, Austria. http://www.r-project.org/.

39. Richard, F.-J., & Hunt, J. H. (2013). Intracolony chemical communication in social insects. Insectes Sociaux, 60(3), 275–291. doi:10.1007/s00040-013-0306-6

40. Roulston, T. H., Buczkowski, G., & Silverman, J. (2003). Nestmate discrimination in ants: Effect of bioassay on aggressive behavior. Insectes Sociaux, 50(2), 151–159. doi:10.1007/s00040-003-0624-1

41. Sanada-Morimura, S., Minai, M., Yokoyama, M., Hirota, T., Satoh, T., & Obara, Y. (2003). Encounter-induced hostility to neighbors in the ant *Pristomyrmex pungens*. Behavioral Ecology, 14(5), 713–718. doi:10.1093/beheco/arg057

42. Sasaki, T., & Tsuji, K. (2003). Behavioral property of unusual large workers in the ant, *Pristomyrmex pungens*. Journal of Ethology, 21(2), 145–151. doi:10.1007/s10164-002-0085-4

43. Satow, S., Satoh, T., & Hirota, T. (2013). Colony fusion in a parthenogenetic ant, Pristomyrmex punctatus. Journal of Insect Science, 13(1), 38. doi:10.1673/031.013.3801

44. Savolainen, R., & Vepsäläinen, K. (2003). Sympatric speciation through intraspecific social parasitism. Proceedings of the National Academy of Sciences, 100(12), 7169–7174. doi:10.1073/pnas.1036825100

45. Sturgis, S., & Gordon, D. (2012). Nestmate recognition in ants (Hymenoptera: Formicidae): a review. Myrmecological News, 16, 101–110.

46. Suarez, A. V., Tsutsui, N. D., Holway, D. A., & Case, T. J. (1999). Behavioral and genetic differentiation between native and introduced populations of the Argentine ant. Biological Invasions, 1, 43–53. doi:10.1023/A:1010038413690

47. Tsuji, K. (1988a). Inter-colonial incompatibility and aggressive interactions in *Pristomyrmex pungens* (Hymenoptera: Formicidae). Journal of Ethology, 6(2), 77–81. doi:10.1007/BF02350871

48. Tsuji, K. (1988b). Obligate parthenogenesis and reproductive division of labor in the Japanese queenless ant *Pristomyrmex pungens*: comparison of intranidal and extranidal workers. Behavioral Ecology and Sociobiology, 23(4), 247–255. doi:10.1007/BF00302947

49. Tsuji, K. (1990a). Kin recognition in *Pristomyrmex pungens* (Hymenoptera: Formicidae): asymmetrical change in acceptance and rejection due to odour transfer. Animal Behaviour, 40(2), 306–312. doi:10.1016/S0003-3472(05)80925-X

50. Tsuji, K. (1990b). Reproductive division of labour related to age in the Japanese queenless ant, *Pristomyrmex pungens*. Animal Behaviour, 39(5), 843–849. doi:10.1016/s0003-3472(05)80948-0

51. Tsuji, K. (1994). Inter-colonial selection for the maintenance of cooperative breeding in the ant *Pristomyrmex pungens*: a laboratory experiment. Behavioral Ecology and Sociobiology, 35, 109–113. doi:10.1007/BF00171500.

52. Tsuji, K., & Dobata, S. (2011). Social cancer and the biology of the clonal ant *Pristomyrmex punctatus* (Hymenoptera: Formicidae). Myrmecological News, 15, 91–99.

53. van Zweden, J. S., & d’Ettorre, P. (2010). Nestmate recognition in social insects and the role of hydrocarbons. Pp. 222–243 in: Bagnères, A.-G. & Blomquist, G. J. (eds.), Insect Hydrocarbons: Biology, Biochemistry, and Chemical Ecology. Cambridge University Press. doi:10.1017/CBO9780511711909.012

54. Wilson, E. O. (1971). The Insect Societies. Belknap Press of Harvard University Press.

55. Wilson, E. O. (1975). Sociobiology: The New Synthesis. Belknap Press of Harvard U Press.

56. Yagound, B., Dogantzis, K. A., Zayed, A., Lim, J., Broekhuyse, P., Remnant, E. J., Beekman, M., Allsopp, M. H., Aamidor, S. E., Dim, O., Buchmann, G., & Oldroyd, B. P. (2020). A single gene causes thelytokous parthenogenesis, the defining feature of the Cape honeybee *Apis mellifera capensis*. Current Biology, 30(12), 2248–2259. doi:10.1016/j.cub.2020.04.033

