## Supporting Information for "Discrimination against non-nestmates functions to exclude socially parasitic conspecifics in an ant"

**Table S1**. Source colonies used in this study.

| ID | Cheaters | Latitude (°N) | Longitude (°E) |
|---|---|---|---|
| A | absent | 34.192433 | 136.32142 |
| B | absent | 34.192186 | 136.321749 |
| C | absent | 34.179448 | 136.294996 |
| D | absent | 34.179519 | 136.29472 |
| F | absent | 34.182122 | 136.293588 |
| H | present | 34.182168 | 136.293528 |
| I | present | 34.181357 | 136.294318 |
| J | present | 34.159249 | 136.274596 |
| K | present | 34.180364 | 136.311384 |
| M | present | 34.183902 | 136.309079 |

**Table S2**. Combination of source colonies in the colony introduction assay.

| Intruder source \ Host source | Type | A | B | C | D | F |
|---|---|---|---|---|---|---|
| H | Cheater | ● | ● | ● | | |
| | Forager | ● | ● | ● | | |
| | Nurse | ●○○ | ● | ●○ | | |
| I | Cheater | | ● | ● | ● | |
| | Forager | | ● | ● | ● | |
| | Nurse | | ● | ● | ● | |
| J | Cheater | | | ● | ● | ● |
| | Forager | | | ●○ | ● | ● |
| | Nurse | | | ● | ●○ | ● |
| K | Cheater | ● | | | ● | ● |
| | Forager | ● | | | ● | ● |
| | Nurse | ●○○ | | | ● | ● |
| M | Cheater | ● | ● | | | ● |
| | Forager | ●○ | ● | | | ● |
| | Nurse | ●○○ | ● | | | ●○ |

| Host source \ Host source | Type | A | B | C | D | F |
|---|---|---|---|---|---|---|
| A | Nestmate | ●●● | | | | |
| B | Nestmate | | ●●● | | | |
| C | Nestmate | | | ●●● | | |
| D | Nestmate | | | | ●●● | |
| F | Nestmate | | | | | ●●● |

For each combination of intruder source and host source colonies, filled circles indicate the trials used for both the behavioral analysis (3.1) and the analysis of the fate of the intruders (3.2), while open circles indicate the trials used only for the behavioural analysis.

**Figure S1**. Behabioural correspondence between the focal intruder and host workers.

For each intruder type, pairs of behavioural scores observed on encounters were stacked across trials into an alluvial plot. Thin translucent curves connect the corresponding scores. The correspondences shown by light orange curves were analyzed in the main text.

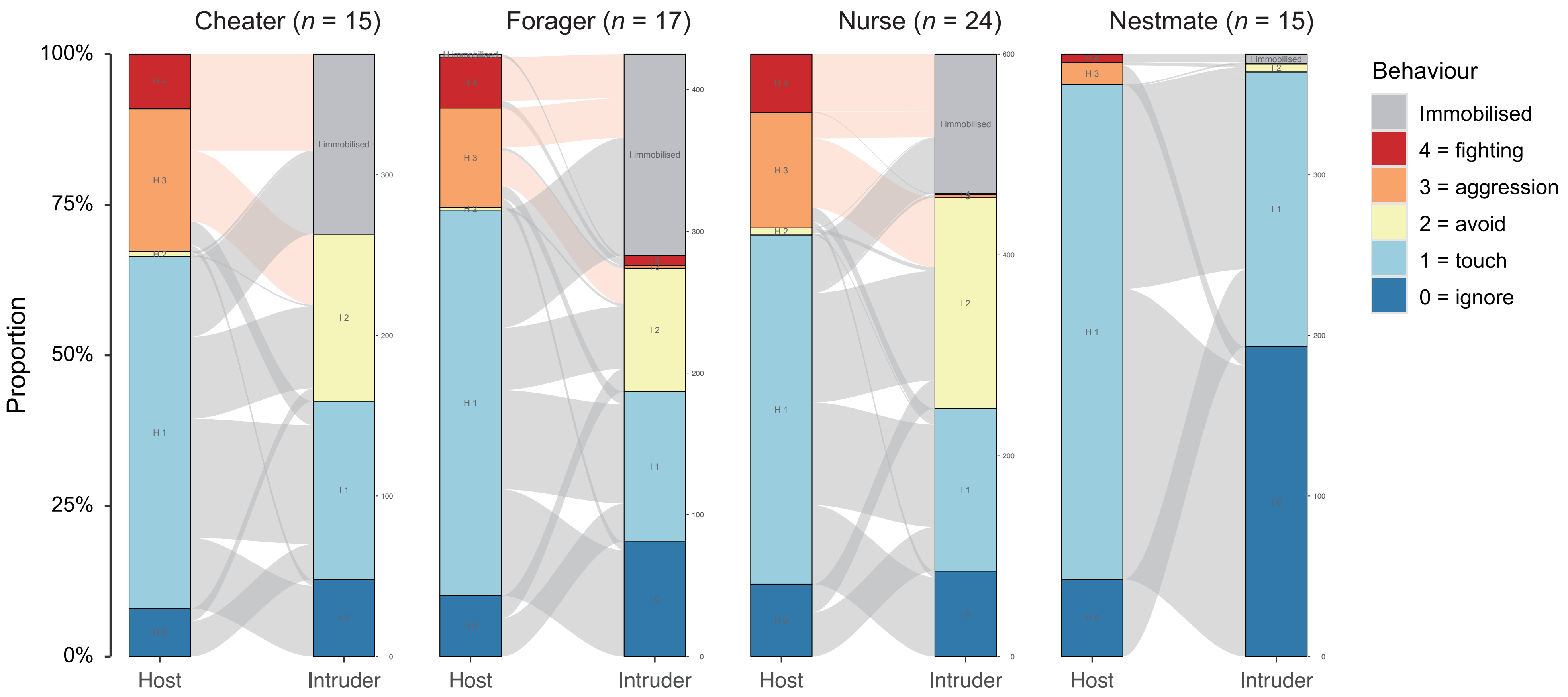
